# Addressable, “Packet-Based” Intercellular Communication through Plasmid Conjugation

**DOI:** 10.1101/591552

**Authors:** John P. Marken, Richard M. Murray

## Abstract

We develop a system for implementing “packet-based” intercellular communication in an engineered bacterial population via conjugation. Our system uses gRNA-based identification markers that allow messages to be addressed to specific strains via Cas9-mediated cleavage of messages sent to the wrong recipient, which we show reduces plasmid transfer by four orders of magnitude. Integrase-mediated editing of the address on the message plasmid allows cells to dynamically update the message’s recipients *in vivo*. As a proof-of-concept demonstration of our system, we propose a linear path scheme that would propagate a message sequentially through the strains of a population in a defined order.

## Introduction

One of the major goals of synthetic biology is to engineer multicellular systems that are able to perform complex tasks [1]. Such systems require communication to occur between the engineered cells in the population, and typically quorum sensing has been used as the architecture of choice due to its simplicity and efficiency [2-4]. However, the high crosstalk between different quorum sensing systems makes it difficult to transmit messages with a high information content between cells— to date, there has been no experimental demonstration of the simultaneous use of more than three orthogonal quorum sensing systems [5, 6]. As engineered multicellular systems become more complex, they will need to accommodate information transfer involving messages with high complexity and dimensionality.

This need for high-capacity information transmission between engineered cells motivates the design of a packet-based intercellular communication system. In packet-based communication, a single architecture is used to transmit messages independently of the content of the messages themselves. While the transmission of an *n*-dimensional message in a quorum sensing system would require *n* orthogonal quorum channels, a single packet-based communication channel would ideally be able to transmit messages of arbitrary length and complexity.

Plasmid conjugation is a naturally-occurring form of horizontal gene transfer that can be used as the basis for a biological “packet-based” intercellular communication system. A conjugative plasmid encodes the genes required to transfer itself to a neighboring cell, collectively called the mobilization elements. Transfer is initiated by the binding of the relaxosome complex to the origin of transfer (*oriT*) on the conjugative plasmid and continues with the replication of the plasmid such that one strand is transferred through a pilus from the sender cell into the receiver cell. There, the plasmid either recircularizes or integrates itself into the receiver cell’s genome [7, 8]. Synthetic conjugation systems have been developed where the *oriT* sequence is mutated or removed from a conjugative plasmid to create a helper plasmid that can no longer transfer itself but, because it still encodes the mobilization elements, confers on its host the ability to transfer any other plasmid that contains the *oriT* sequence [9, 10].

A “packet-based” intercellular communication framework based on such a synthetic conjugation system would have the following desirable properties: (1) because the helper plasmid cannot transfer itself, one can engineer explicit distinctions between senders and receivers within a population; (2) because the message is encoded into the sequence of the plasmid, the system can easily transmit messages with high information content and dimensionality; (3) one can choose from a number of information-encoding schemes, for example choosing to represent a message in either DNA, RNA, or protein expression; (4) conjugation occurs successfully for plasmids at least as large as 100 kb [11] and at least as small as 10 kb [12], meaning that the system can indeed act as a “packet-based” communication system where the mechanism of transmission is unaffected by the length or content of the message.

Below, we describe a design for a “packet-based” intercellular communication system that, in addition to having the above properties, allows messages to be addressed so that they can only be transferred to a defined subset of the strains within a population.

### Addressable Information Transfer

The basis for message addressability in our system is that each strain expresses Cas9 and a unique guide RNA (gRNA). An ‘address’ region can then be encoded onto the message plasmid that contains binding sites for the gRNAs associated with certain strains. If a plasmid is sent to a cell whose gRNA matches a site on the address region, then the Cas9 in the receiver cell will cleave and degrade the plasmid, blocking the receipt of the message. With this system, one can ensure addressability to any subset of an *n*-strain population simply by expressing *n* orthogonal gRNAs. As each gRNA binding site is only 20bp in length, the address region has the additional advantage of being compact.

The fact that the address region is encoded on the message plasmids also allows the cells themselves to alter its content. By flanking the gRNA binding sites in the address region with integrase attachment sites, it is possible for the cells to express the corresponding integrases to modularly insert, remove, and swap binding sites from the address region, dynamically updating the recipient list for their messages [13]. The expression of these integrases could be regulated by signaling within the population circuit itself, enabling feedback-driven autonomous regulation of information flow within a population. As the length of an integrase attachment site typically ranges between 60 - 70 bp [14], the ability to edit the address region does not significantly add to its length. Full modularity in address modification can be obtained with *n* orthogonal integrases for an *n*-strain population— currently there are at least 11 known orthogonal integrase/excisionase pairs [14].

## Results

### Steady-state transconjugant cell density is not strongly affected by the length of coculturing time and the initial density of senders and receivers

We first wanted to assess the efficiency of plasmid transfer, to determine if the system is viable as a basis for intercellular signaling. We cocultured sender cells (containing the tetracycline (Tet) resistant F_HR plasmid from [9] and a signal plasmid that expresses sfYFP constitutively on a chloramphenicol (Chlor) resistant backbone) with receiver cells (which also contain the F_HR plasmid, as well as a constitutive mScarlet-I on a kanamycin (Kan) resistant backbone). The density of senders, receivers, and transconjugants within the culture could be determined by selective plating— YFP-positive cells on Chlor+Tet plates are Senders, YFP-negative cells on Kan plates are Receivers, and YFP-positive cells on Chlor+Kan plates (as well as YFP-positive cells on Kan plates) are Transconjugants.

Given that previous work has demonstrated that the efficiency of conjugation drops dramatically in stationary phase [15, 16], we expected to find that the density of transconjugants in the culture would be correlated with the length of time that senders and receivers were cocultured in exponential phase— in other words, if senders and receivers were mixed at a low initial density, then we would expect to see more transconjugants at steady state compared to if they were mixed at a high initial density. Interestingly, our data contradicted this expectation— the steady-state transconjugant density appeared to be relatively unaffected by the initial mixing density of the senders and receivers (Fig. 1). These results suggest that the reduction in plasmid transfer rate that occurs in stationary phase can potentially be overcome by other factors, such as a higher density of sender and receiver cells leading to more frequent contacts between cells (and hence more opportunities for conjugation to occur).

**Fig. 1:**
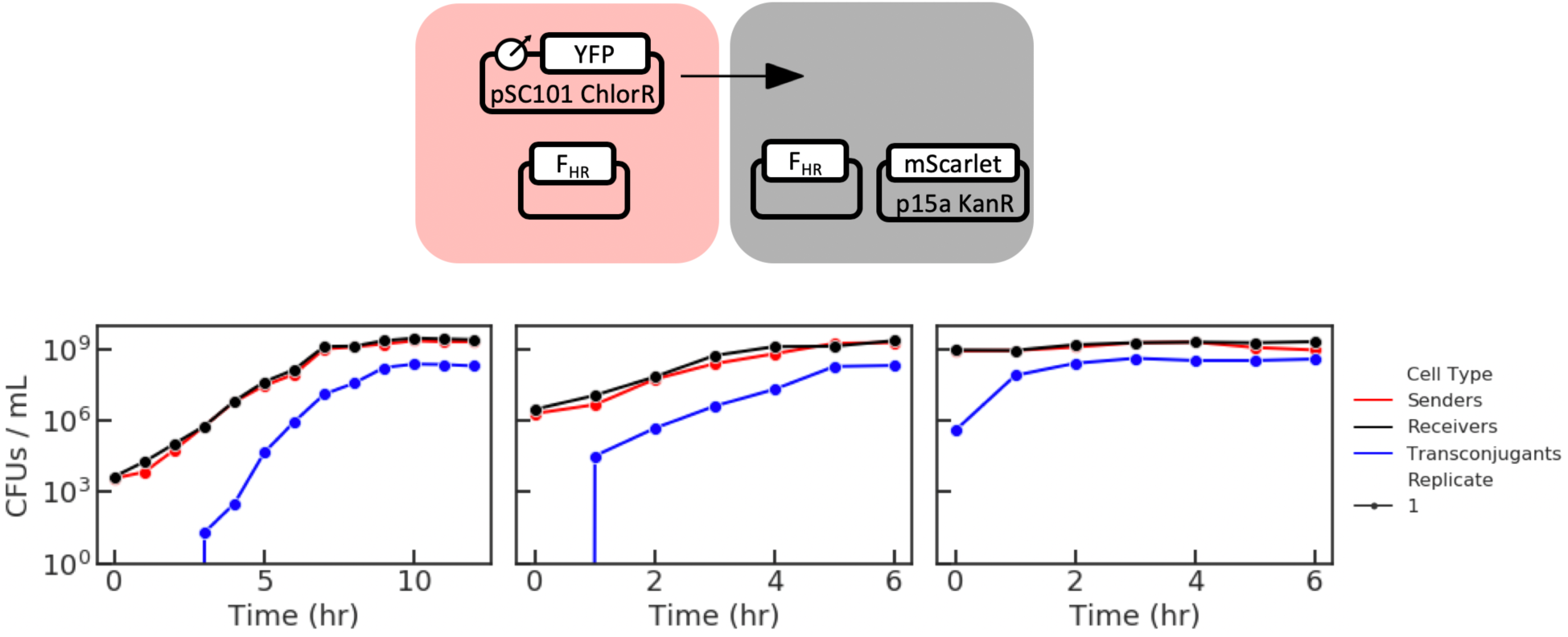
Timecourse measurements of strain densities within a coculture, where senders and receivers were mixed together at different initial cell densities. Top: simplified experimental schematic. The density of transconjugants at the final timepoint were 4%, 5%, and 11% of the total cell density for the low, medium, and high initial density conditions, respectively. Each panel represents a single biological replicate.

### Cas9 can inducibly cut and degrade plasmids *in vivo*

In order for the Cas9 system to successfully block the transfer of message plasmids, it is necessary for Cas9-mediated cleavage of the plasmid at the gRNA binding site to lead to degradation of the plasmid. To determine whether this is the case, we first conducted a fluorescence-based assay for Cas9-mediated plasmid degradation *in vivo*. We designed two LasAHL-inducible sfYFP constructs that each contained a unique gRNA binding site (“Sce” or “U22”) and transformed these plasmids into cells containing a CinAHL-inducible Cas9 construct expressing either the Sce or the U22 gRNA. Cells were induced with variable concentrations of CinAHL to activate Cas9 and were grown for 4 hours to allow the Cas9 time to cleave the target YFP plasmids. The YFP was then induced with 1 uM LasAHL and grown to steady state. Since the gRNA binding site was placed downstream of the YFP transcriptional unit, if Cas9 cleavage only linearized the plasmid rather than leading to its degradation, then in principle YFP could still be expressed from the linear DNA following cleavage. Thus, a lack of fluorescence was interpreted as a proxy for the degradation of the plasmid DNA in the cell.

As expected, cells in which the expressed gRNA matched the binding site on the YFP plasmid displayed little fluorescence, while cells with a mismatch between the gRNA and the binding site on the YFP plasmid were fluorescent (Fig. 2a, left). Importantly, the cleavage of the YFP plasmid by Cas9 does not seem to have a noticeable effect on the maximal density that the cells reach (although increasing induction of Cas9 did decrease the maximal OD700 value in all conditions) (Fig. 2a, right).

**Fig. 2:**
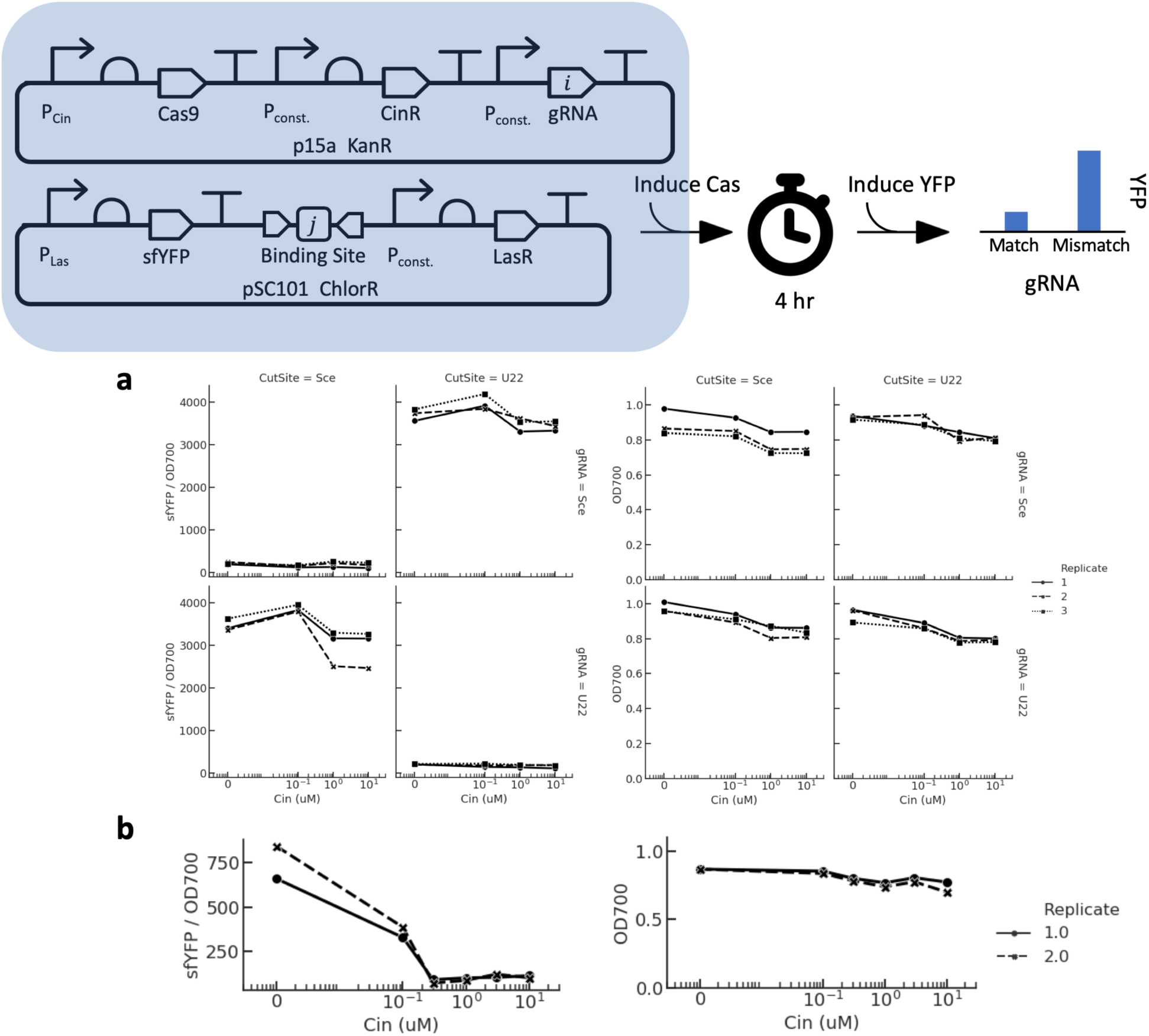
Fluorescence-based assays for Cas9-mediated plasmid degradation. Top: experimental schematic— YFP fluorescence should be low when the gRNA matches the binding site on the YFP plasmid, and should be high otherwise. (a) Cas9-mediated plasmid degradation only occurs when the gRNA matches the binding site on the target plasmid. (b) Using a construct with weaker expression of Cas9 and the Sce gRNA allows inducible degradation of the Sce-site target plasmid. Left: OD-normalized maximal values from fluorescence timecourses. Right: maximal OD values from the same timecourses. All fluorescence measurements are in arbitrary units. Markers/dashes represent different biological replicates (n=3 for (a) and n=2 for (b)).

Although this experiment demonstrated that Cas9-mediated cleave can lead to plasmid degradation *in vivo*, it also demonstrated that the construct we used to express Cas9 was sufficiently leaky such that complete degradation of the YFP plasmid occurred within 4 hours even without the explicit induction of Cas9. As we would like to be able to inducibly control the plasmid cleavage, we altered our Cas9 construct to weaken the expression of the Cas9 and gRNA, and repeated the fluorescence-based plasmid degradation assay for the Sce gRNA vs. Sce binding site condition. With our new Cas9 construct, we were able to observe YFP expression when Cas9 was not induced, as desired (Fig. 2b). However, the fluorescence in the 0 uM CinAHL condition is still lower than the fluorescence reached in the mismatched gRNA conditions (compare Fig. 2b with Fig. 2a), suggesting that there is still an intermediate amount of leaky Cas9-mediated degradation in this weaker construct.

### gRNA binding sites on a plasmid can be swapped *in vivo* to induce or inhibit its degradation

We then investigated whether integrase-mediated cassette exchange could be used to rescue plasmids from Cas9-mediated degradation. We transformed the inducible YFP plasmid containing the U22 binding site flanked by attP sites for the TP901 integrase, along with a plasmid expressing TP901 and containing a Sce binding site flanked by TP901 attB sites, into cells containing a plasmid encoding an inducible Cas9 with either the Sce or the U22 gRNA. In these cells, the expression of the TP901 integrase should result in cassette exchange to swap the U22 binding site on the YFP plasmid with the Sce binding site. The fluorescence-based degradation assay described above was then repeated on these cells, with Cas9 induced with 1 uM CinAHL for 4 hours prior to induction of YFP with 1 uM LasAHL.

As expected, the YFP plasmid was not degraded by the U22 gRNA, as integrase-mediated cassette exchange had swapped the U22 binding site on the YFP plasmid to the Sce site prior to induction of Cas9 (Fig. 3). This swapping was further confirmed by the fact that the YFP plasmid was degraded by the Sce gRNA.

**Fig. 3:**
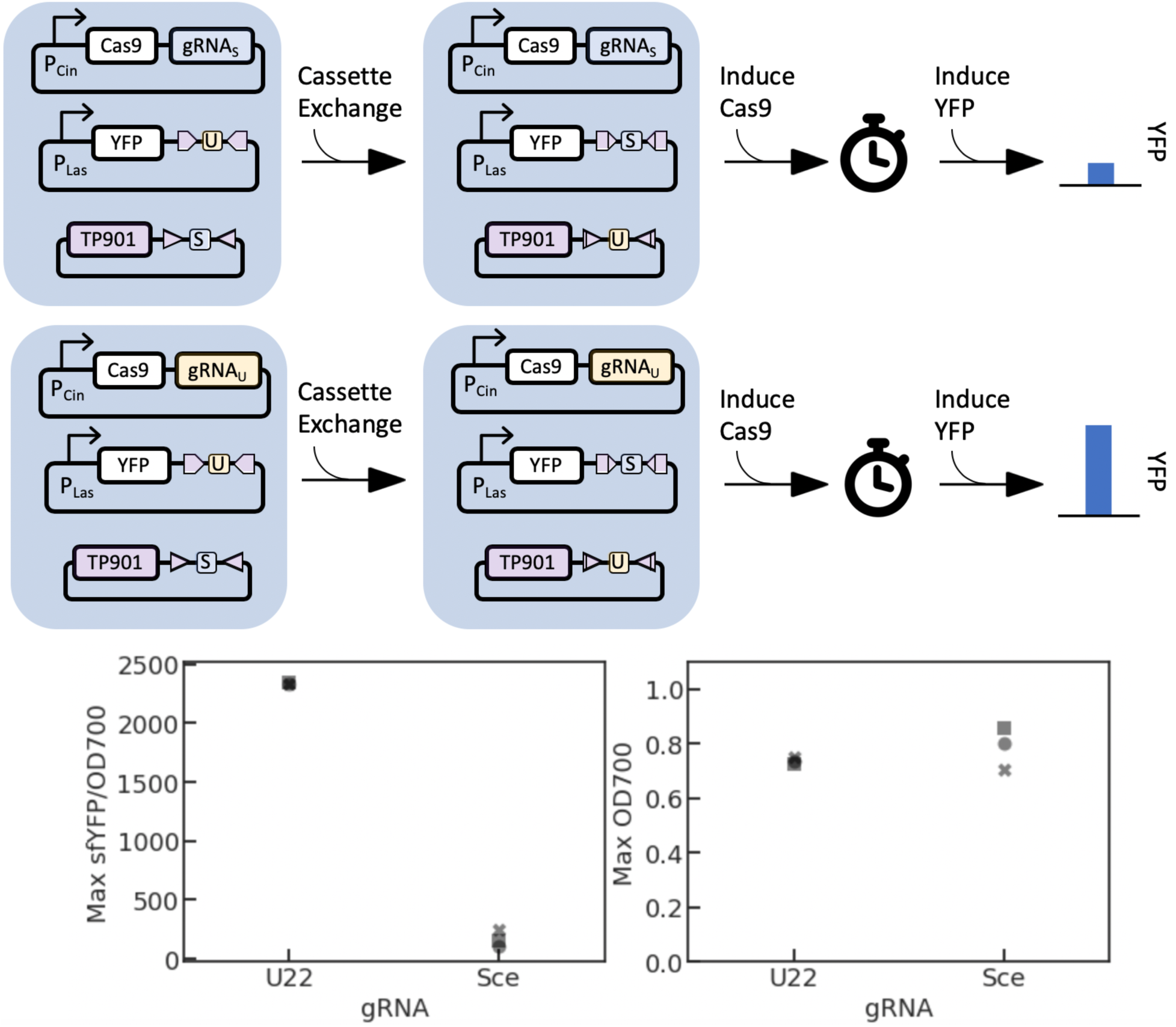
Integrase-mediated cassette exchange rescues target plasmids from Cas9-mediated degradation. Top: simplified experimental schematic— integrase-mediated cassette exchange should swap the U22 binding site on the YFP plasmid for the Sce site, causing the YFP plasmid to be degraded by only the Sce gRNA and not the U22 gRNA. Left: OD-normalized maximal values from fluorescence timecourses. Right: maximal OD values from the same timecourses. All fluorescence measurements are in arbitrary units. Markers represent different biological replicates (n=3).

### Cas9-mediated cleavage can inducibly block the receipt of transferred plasmids

Once we determined that Cas9-mediated plasmid degradation was possible *in vivo*, we then investigated whether the presence of Cas9 and a matching gRNA is sufficient to block the receipt of a plasmid transferred from a sender cell. We cocultured receiver cells, containing the CinAHL-inducible Cas9 construct with the U22 gRNA on a Kan-resistant backbone, with sender cells, containing the F_HR plasmid and a message plasmid constitutively expressing sfYFP on a Chlor-resistant vector containing either the U22 or Sce binding site. Cas9 was induced in the receiver cells with either 0 uM or 1 uM CinAHL for an hour prior to mixing with the sender cells, and the coculture was incubated for 5 hours after mixing.

We found that expression of Cas9 with a matching gRNA in the receiver cells was indeed able to inhibit the receipt of the signal plasmid, with the final density of transconjugants in the matching-gRNA condition being on average 0.0004% of the total cell density when Cas9 was induced, compared to 3.5% when it was not, which is almost a 10,000-fold reduction (Fig. 4, top row). Additionally, we found that leaky expression of Cas9 with a matching gRNA only slightly inhibits the receipt of a message plasmid— the final transconjugant densities were, on average, 8.4% of the total cell density in the mismatched gRNA conditions, suggesting that leaky expression of Cas9 with a matching gRNA only reduces final transconjugant density by approximately 2.4-fold (Fig. 4, middle row).

**Fig. 4:**
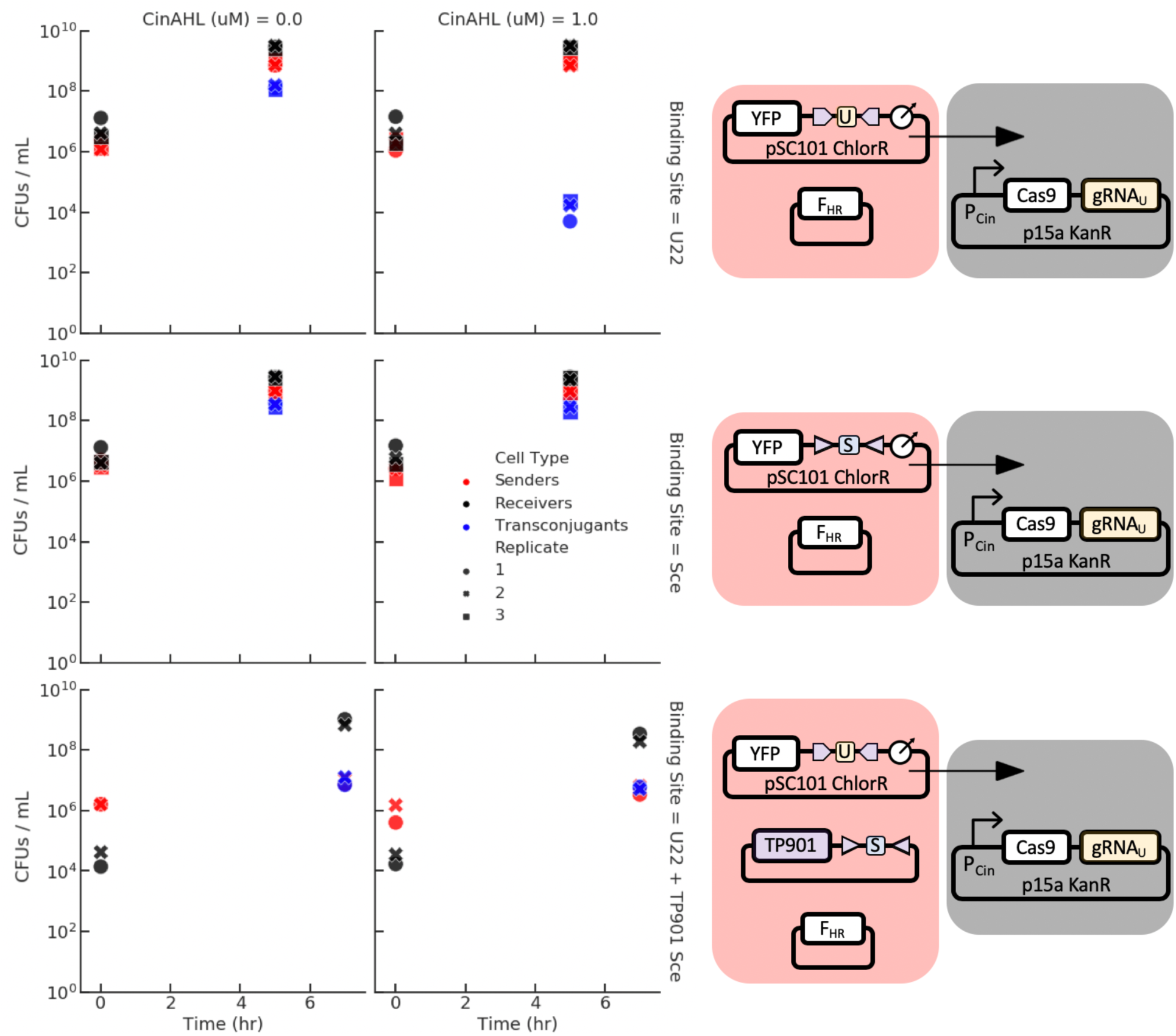
Cas9-mediated plasmid cleavage inducibly blocks plasmid transfer if and only if the gRNA in the receiver matches the binding site on the message plasmid. Right: simplified experimental schematic— when Cas9 with U22 gRNA is induced by CinAHL, the transfer of plasmids with the U22 site should be blocked, while the transfer of plasmids with the Sce site should be unaffected. Bottom row: integrase-mediated cassette exchange should swap the U22 site on the message plasmid for a Sce site, allowing it to transfer into receiver cells expressing Cas9 and the U22 gRNA. Different markers styles represent different biological replicates (n=3 top and middle rows, n=2 bottom row).

### Integrase-mediated editing of the address region can alter the recipients of a signal plasmid

We then repeated the above experiment but including integrase-mediated editing of the address region. Our sender cells now contained F_HR and a constitutive YFP message plasmid with the U22 gRNA binding site (flanked by TP901 attP sites) on a Chlor-resistant backbone, as before, but also a constitutively-expressed TP901 with a Sce gRNA binding site flanked by TP901 attB sites on a Kan-resistant backbone. Our receiver cells now contained the CinAHL-inducible Cas9 vector with the U22 gRNA on a Kan-resistant backbone, as before, and additionally a carbenicillin (Carb) resistant vector. Senders were counted as YFP-positive cells on Chlor+Kan plates, Receivers were counted as YFP-negative cells on Kan+Carb plates, and Transconjugants were designated as YFP-positive cells on Chlor+Carb plates. As before, Cas9 was induced with either 0 uM or 1 uM CinAHL for an hour prior to mixing.

As expected, we saw that the presence of the integrase in the sender cells rescued the message plasmid from blockage by the U22 gRNA, although there was a notable decrease in the transfer efficiency compared to the earlier transfer experiments without TP901 (Fig. 4 bottom row vs. top and middle rows). This is likely due to the slow growth of the Sender strain observed in the cassette exchange experiments. Nonetheless, we observed that the transfer efficiency was essentially unaffected by the presence of Cas9 (transconjugants were, on average, 2% of the population with 1 uM CinAHL compared to 1% without CinAHL), which further supports the fact that TP901 was able to fully edit the address regions in the message plasmids.

## Discussion

Our addressable intercellular communication system allows one to engineer the flow of information through a population. For example, one could design a linear path scheme where a message must propagate sequentially through the strains in a population in a defined order, without backtracking or skipping ahead. A three-node linear path can be implemented using only two gRNAs and one integrase (Fig. 5a)— for a message traveling from strain A to B to C, its address initially contains the binding site for the C gRNA, preventing direct transfer of the message from strain A to strain C. Strain B expresses an integrase and a cassette containing the binding site for the A gRNA, such that once the message plasmid is sent to a cell in the B strain, integrase-mediated cassette exchange [13] swaps the binding site on the message plasmid from the C binding site to the A binding site. Once this occurs, the message plasmid can no longer return to strain A, but can now be sent to strain C. The linear path is generalizable to any *n*-strain population and can be implemented using *n-1* orthogonal gRNAs and *n-2* orthogonal integrases for *n* ≥ 4 (Fig. 5b). This concept of information routing can be extended to more complicated paths that could be dynamically updated by the population circuit itself.

**Fig. 5:**
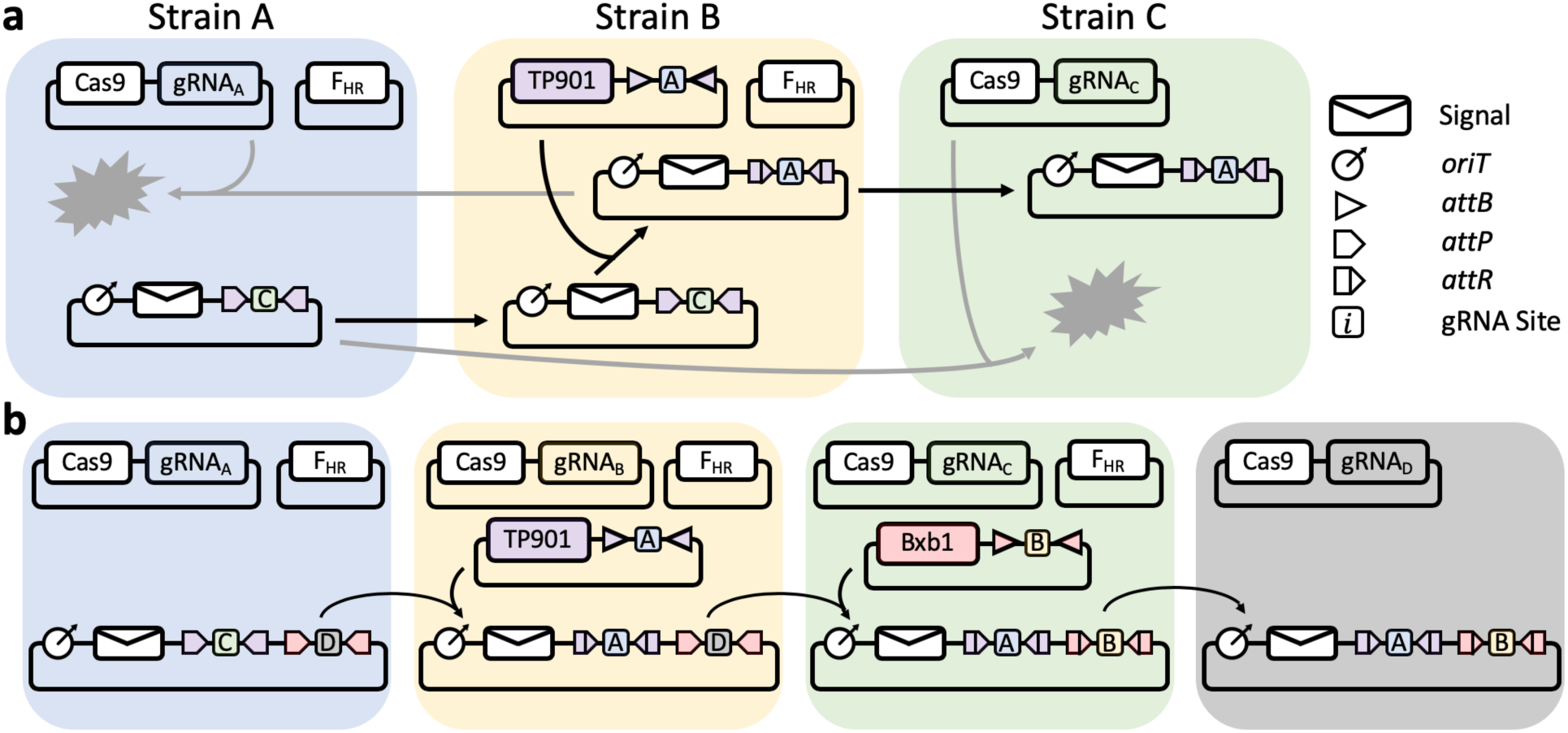
A linear-path scheme for routing a message through a population. **(a)** A 3-strain linear path created with one integrase (TP901) and two gRNAs (“A”,”C”). The terminal recipient (strain C) does not contain a helper plasmid, i.e. cannot send signals so a gRNA for strain B is not required. **(b)** A 4-strain linear path created with two integrases (TP901, Bxb1) and four gRNAs.

Although previous work has explored the use of intercellular DNA transfer as the basis for a synthetic communication system [17-19], including the 2006 Berkeley iGEM team that used riboswitch lock/key pairs to implement message addressability by having only specific receiver strains properly interpret a promiscuously transmitted message [20], we believe our system is the first to experimentally implement the ability to selectively transmit messages to a specific subset of a population. We hope that our work can serve as the basis for future work that continues to incorporate the specific advantages of packet-based communication into intercellular information transfer systems.

In addition to serving as the basis for an intercellular packet-based communication system, plasmid conjugation has the additional advantage of occurring only between spatially-adjacent cells [21]. This provides a potential contact-based signaling mechanism for the prokaryotic synthetic toolbox, which could enable the design of spatial patterning circuits with greater resolution than is currently feasible using diffusible signaling molecules.

Although the use of plasmid conjugation-based signaling may not be as fast or efficient as quorum-sensing based signaling, it is important to have a variety of signaling architectures at synthetic biologists’ disposal. This way, different architectures can be designed into multicellular systems in such a way that different requirements in the system can be addressed by the specific advantages of different signaling architectures. As engineered multicellular systems become more complex, the need for such a diverse toolbox of signaling architectures will become increasingly more important. This work provides an element in this toolbox that has specific advantages in message addressability and high-dimensional information transfer.

## Materials and Methods

### Strains and Constructs

All cells containing the F_HR plasmid are MG1655 *E. coli* cells, while all other cells used in this study are JS006 strain *E. coli*.

All constructs were assembled using 3G assembly [22].

Unless otherwise noted, antibiotic selection was always applied to media such that all plasmids in all strains in a culture were under selection. Antibiotic concentrations used were: 100 mg/mL for carbenicillin, 34 mg/mL for chloramphenicol, 50 mg/mL for kanamycin, and 20 mg/mL for tetracycline. These concentrations were also used for antibiotic selection plates. When minimal media was used, these antibiotic concentrations were halved.

### Coculturing Assays

Sender and receiver cells were grown from a single colony overnight in 3 mL of 2xYT media in a 15 mL culture tube, then diluted 1:100 into 3 mL fresh 2xYT media in the morning. Cells were then grown for approximately 2-3 hours until midlog— in Cas9 blocking assays, receiver cells were diluted 1:2 into fresh 2xYT media with either 0 or 1 uM CinAHL after the first hour of growth. Cells were then spun for 5 minutes at 3,500 rpm in an Eppendorf 5810 R ultracentrifuge and resuspended in 3 mL of fresh 2xYT media with no antibiotics. 50 uL of sender cell culture and 50 uL of receiver cell culture were then mixed into 5 mL of fresh 2xYT media with no antibiotics in a 250 mL flask and incubated at 37C and 230 rpm shaking. For stationary phase transfer assays, the overnight cultures were spun down and resuspended in no-antibiotic media and mixed by adding 2.5 mL of the sender cell culture to 2.5 mL of the receiver cell culture. Selective plating onto 5 mL LB agar plates was performed with antibiotic selection combinations as described in the main text. The first timepoint was plated immediately after mixing the sender and receiver cells. Serial dilutions of the coculture were performed by transferring 10 uL of culture to 990 uL of fresh 2xYT media with no antibiotics to do a 100-fold dilution. 100 uL of culture was plated onto each plate.

Colony counting and fluorescence determination was performed by imaging each plate on an Olympus MVX10 microscope using the YFP filter from a Sutter 10-2 fluorescence filter wheel and manually counting the images in ImageJ.

### Fluorescence Assays

Cells were grown from a single colony overnight in 3 mL of M9CA minimal media (Teknova). Cells were diluted 1:100 into 300 uL fresh M9CA media without chloramphenicol in the morning in a 96-well square-well plate (Brooks MGB096-1-2-LG-L) and grown at 37C with maximum-speed linear shaking in a BioTek Synergy H1m. YFP fluorescence was measured with excitation/emission at 503/540 nm. Optical Density was determined by measuring the absorbance of the cultures at 700 nm.

For Cas9-mediated degradation assays, Cas9 was induced with varying levels of CinAHL once the OD700 of the cultures in each well were around 0.2. Induction was performed alongside a 1:10 dilution by removing 270 uL of media from each the well and adding 270 uL of fresh M9CA media (without chloramphenicol) containing a concentration of inducer calibrated to reach the target concentration in the full 300 uL volume. After 4 hours, YFP was induced with 1 uM LasAHL (while still preserving the CinAHL concentration) alongside a 1:10 dilution in the same way.

For assays of integrase-mediated rescue from Cas9-mediated degradation, the same induction procedures were performed as described above, but beginning with 0 or 10 uM of salicylate induction of TP901 (into M9CA with the full antibiotic profile) for 4 hours prior to the induction of Cas9 and YFP as described above.

## Acknowledgments

We thank Andrey Shur and Andy Halleran for insightful discussions. The F_HR plasmid used in [9] was generously provided by Tatiana Dimitriu. PSal, PLas, and PCin promoters, as well as their respective activators NahR, LasR, and CinR, were a generous gift from Adam Meyer (now documented in [23]). Plasmid vectors were provided by Douglas Densmore at the Cross-disciplinary Integration of Design Automation Research lab (Addgene Kit # 1000000059). This research is supported by the Institute for Collaborative Biotechnologies through cooperative agreement W911NF-19-2-0026 and grant W911NF-09-0001 from the U.S. Army Research Office. The content of the information on this page does not necessarily reflect the position or the policy of the Government, and no official endorsement should be inferred.

